# Effect of Chlonisol (2-[3-(2-chloroethyl)-3-nitrosoureido]-1,3-propanediol) on Overall Survival in Laboratory Rodents with Intracranial Tumors: A Meta-analysis of Preclinical Studies

**DOI:** 10.1101/2022.04.14.488286

**Authors:** Iaroslav G. Murazov, Alexander N. Stukov, Iuliia G. Zmitrichenko

## Abstract

**Background:** The need for new, effective, and affordable drugs for the treatment of primary and metastatic brain tumors remains unsatisfactory. Preclinical studies of chlonisol (2-[3-(2-chloroethyl)-3-nitrosoureido]-1,3-propanediol) showed promising results in the treatment of experimental intracranial tumors.

**Objectives:** To apply a meta-analytical approach to estimate the combined effect size of chlonisol on overall survival (OS) in rodents with brain tumor transplants.

**Search strategy:** Data for the meta-analysis were obtained from the laboratory’s internal database from reports of preclinical studies of chlonisol.

**Selection criteria:** Eligible studies were parallel preclinical trials in rodents (mice, rats) with intracranially transplanted tumors. Chlonisol was compared with active control treatment (lomustine or temozolomide). All cytostatics were administered at the maximum tolerated dose (MTD). The duration of the studies was at least 90 days. The main outcome was OS-HR (hazard ratio).

**Data analysis:** We applied the inverse variance technique for the meta-analysis of HRs. In HR analysis we adopted a random-effect model.

**Results:** The analysis included seven trials with 132 rodents. Studies were conducted between 2016 and 2022. As a murine intracranial grafts we used Ehrlich’s carcinoma, Sarcoma 180 and the HER2-positive mammary tumor derived from a female FVB/N HER-2/neu transgenic mouse. Glioma 35 was transplanted into rats. Compared with active control, oral or intraperitoneal administration of chlonisol at MTD of 20 mg/kg, significantly reduced the risk of death by 63% (HR=0.37; 95% CI: 0.24-0.56; P<0.00001) in animals with intracranial tumors. The direction in favor of chlonisol was stable across studies despite the use of different animals and transplants, the routes of administration of chlonisol, and control treatment. No significant heterogeneity was observed between the studies (Tau^2^ = 0.03; Chi^2^ = 6.52; df = 6; P = 0.33; I^2^ = 8%).

**Conclusion:** Compared with lomustine and temozolomide, chlonisol treatment in MTD provides an important advantage in OS in animals with intracranial tumors. Our results may serve as a basis for further study of chlonisol as a chemotherapy agent for primary and metastatic brain tumors.

## INTRODUCTION

Primary brain tumors (BT) in adults are rare and account for about 3% of all types of malignant neoplasms. BT, however, causes the greatest loss of years of life in all types of cancer (Bates *et al*. 2018). BTs are the most common type of solid tumor in children and the leading cause of cancer death in this population (Jones *et al*. 2019). Brain metastases are 10 times more common than primary tumors (Vargo *et al*. 2017). The most common metastases of the central nervous system (CNS) occur in lung tumors, breast tumors, and malignant tumors [Ostrom *et al*. 2018]. The overall survival (OS) of patients with primary BT has not changed much over the past 10 years, despite the introduction of new methods of treatment and advances in understanding the growth biology of tumors of this localization. The search for effective and affordable drugs for the treatment of primary BTs and brain metastases remains an urgent task of modern oncopharmacology.

The object of the presented study is the domestic antitumor compound (2-[3-(2-chloroethyl)-3-nitrosoureido]-1,3-propanediol) (chlonisol). Chlonisol belongs to the class of nitrosoalkylureas (NAU) (Fig. 1). The molecular mechanisms of the action of NAU are associated with the alkylation and carbamoylation of macromolecules by the products of the biodegradation of the compounds. As far as cytogenetic activity is concerned, chlonisol exceeds all other NAU-class drugs studied earlier. (Stukov *et al*. 2020).

**Figure 1.**
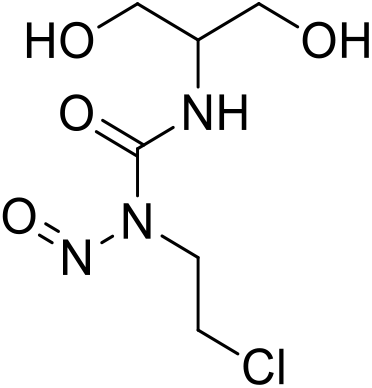
2-[3-(2-chloroethyl)-3-nitrosoureido] -1,3-propanediol (chlonisol).

Meta-analysis (MA) is a statistical analysis of the results of several comparable studies on the same problem (generally treatment, prevention, diagnosis, etc.). The combination of studies offers a larger sample for analysis and greater statistical power. In health care, MA has been used actively in recent decades to critically assess the results of clinical trials. MA enables doctors and patients to make informed decisions on medical services and the implementation of specific interventions (Ioannidis *et al*. 2000). The idea of generalized statistical estimation of preclinical research is relatively new. However, the goals and objectives of meta-analysis of preclinical studies differ fundamentally from those of meta-analysis of clinical trials results (Vesterinen *et al*. 2014). First, systematic reviews and meta-analysis of preclinical studies are hypothesis generating and pursue exploratory goals, allowing to include all available data on a topic of interest. Secondly, meta-analysis of preclinical studies allows for greater translation of results into subsequent clinical studies and ensures the objectivity of the results while observing the principles of 3 R’s (replacement, reduction, refinement). This statistical tool also reduces the total cost of preclinical studies. Thirdly, an important task of meta-analysis of preclinical studies is to identify sources of possible bias in order to continue to prepare for the design of clinical protocols and improve the quality of its implementation. Finally, for a meta-analysis of preclinical studies, an overall estimate of the effect size of an intervention is not entirely important. More important is the assessment of the direction of the effect, its «stability» and the identification of potential sources of heterogeneity between studies (Hooijmans *et al*. 2014).

The objective of the meta-analysis in this study is to provide the size of the combined effect of chlonisol on OS in mice and rats with intracranial tumor transplants, compared with known chemotherapeutic drugs.

## MATERIALS AND METHODS

### Search strategy and selection of studies. Inclusion criteria

Data for meta-analysis were obtained from the internal database of the laboratory of cancer chemoprevention and oncopharmacology of the FSBI «N.N. Petrov National Medical Research Centre of Oncology» of the Ministry of Healthcare of the Russian Federation. We used the following criteria to include studies in meta-analysis:

1. Study of the antitumor activity of clonisol in mice or rats.
2. Intracranial transplantation of syngeneic or lineage-nonspecific allografts according to the method described by Murazov *et al*. (2021). Briefly, in the animal facility operating room, animals were inoculated under isoflurane anesthesia (induction - 3%, maintenance - 1.5%) with a shortened to 3 mm and newly sharpened injection needle (23 G) into the supraorbital region near the midline of the head, through which a suspension of tumor cells was inoculated in a sterile 0.9% normal saline.
3. The presence of the comparison group with the active drug.
4. The study lasted at least 90 days.
5. As an endpoint, the study included an assessment of OS.

All studies were approved by the local ethics committee and carried out in accordance with the European Convention for the Protection of Vertebrate Animals Used for Experimental and Other Scientific Purposes: Appendix A of the ETS 123 and Directive 2010/63/EU on the protection of animals used for scientific purposes.

### Statistical analysis

The statistical data processing was done using Review Manager (RevMan), version 5.4.1 (Cochrane Collaboration 2020). OS results were expressed as the hazard ratio (HR) and 95% confidence interval (95% CI). For meta-analysis, we chose a random effects model using the inverse variance method, since different animal species, strains/stocks of mice were used in the studies, as well as reference drugs. Between the studies, the route of chlonisol administration and the comparator also differed. The results of the meta-analysis are presented as a forest plot. The statistical heterogeneity was evaluated using the Q-test and the index of heterogeneity I^2^.

## RESULTS AND DISCUSSION

### Search results

A total of 7 studies were found in the laboratory database that assessed the OS of animals treated with chlonisol and compared drugs - lomustine and temozolomide. Preclinical studies were conducted between 2016 and 2022. The results of these studies have been processed by statistical analysis. A total of 132 animals were included. The following intracranial transplants were used in mice: Ehrlich’s carcinoma, sarcoma 180, HER2-positive mammary tumor of the FVB/N HER-2/neu transgenic female mouse. Rats were transplanted intracranially with glioma 35 (Table 1). OS was defined as the time from randomization to animal death from any cause.

**Table 1.** Characteristics of included preclinical studies.

| **Animal species, strain/stock, gender** | **Intracranial transplant** | **Number of animals in exp/control group, N** | **Dose regimen and route of administration of chlonisol** | **Dose regimen and route of administration of control treatment** | **Median OS exp vs control, days** | **HR (95% CI); logrank-test P** |
| --- | --- | --- | --- | --- | --- | --- |
| SHR mouse, females | Ehrlich's carcinoma | 10/10 | 20 mg/kg per os single dose 24 h after transplantation | Lomustine 50 mg/kg per os single dose 24 h after transplantation | 15,5 vs 11,5 | 0,24 (0,08-0,73); P <0,0001 |
| BALB/c mouse, males | Ehrlich's carcinoma | 11/11 | 20 mg/kg IP single dose 24 h after transplantation | Lomustine 50 mg/kg per os single dose 24 h after transplantation | 16,0 vs 12,0 | 0,26 (0,09-0,72); P <0,0001 |
| 178/SN mouse, males | Ehrlich's carcinoma | 8/8 | 20 mg/kg IP single dose 72 h after transplantation | Temozolomide 50 mg/kg per os for 5 days, the first dose - 72 h after transplantation | 19,0 vs 16,0 | 0,40 (0,13-1,18); P=0,0222 |
| BALB/c mouse, females | Sarcoma 180 | 10/10 | 20 mg/kg IP single dose 24 h after transplantation | Lomustine 50 mg/kg IP single dose 24 h after transplantation | 26,0 vs 13,0 | 0,16 (0,05-0,54); P <0,0001 |
| FVB/N mouse, males | HER2-positive mammary tumor of the FVB/N HER-2/neu transgenic female mouse | 10/10 | 20 mg/kg IP single dose 72 h after transplantation | Temozolomide 50 mg/kg per os for 5 days, the first dose - 72 h after transplantation | 55,0 vs 43,0 | 0,36 (0,13-0,99); P=0,0175 |
| FVB/N mouse, males | HER2-positive mammary tumor of the FVB/N HER-2/neu transgenic female mouse | 9/9 | 20 mg/kg IP single dose 72 h after transplantation | Lomustine 50 mg/kg per os single dose 72 h after transplantation | 40,0 vs 31,0 | 0,89 (0,35-2,25); P=0,7848 |
| Wistar rats,<br>males | Glioma 35 | 8/8 | 20 mg/kg IP single dose 24 h<br>after transplantation | Lomustine<br>50 mg/kg IP single dose 24 h after<br>transplantation | 71,0 vs 45,5 | 0,48 (0,14-1,69);<br>P=0,2425 |

**Notes:** IP – intraperitoneal

### OS analysis

The overall effect estimate showed that compared with control treatment oral or intraperitoneal (IP) administration of chlonisol at the maximum tolerated dose (MTD) of 20 mg/kg significantly reduced the risk of death in animals with intracranial tumors by 63% (HR=0.37; 95% CI: 0.24-0.56; P<0.00001) (Fig. 2). The direction of the effect in favor of chlonisol was consistent throughout the studies, despite the use of different animals and transplanted tumors, the routes of administration of chlonisol and reference drugs. No significant between-study heterogeneity was observed (Tau^2^ = 0.03; Chi^2^ = 6.52; df = 6; P = 0.33; I^2^ = 8%).

**Figure 2.**
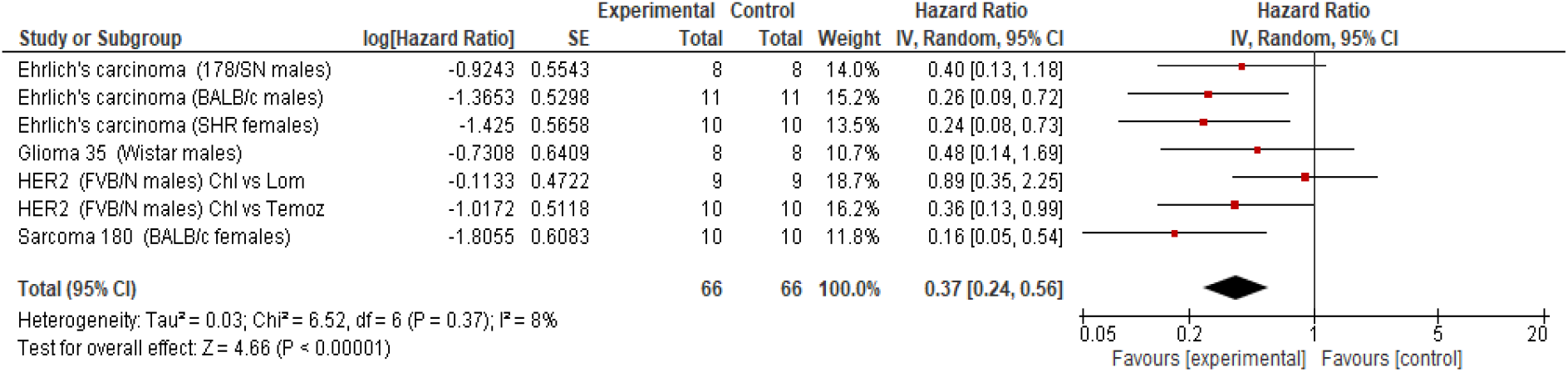
Random effects model for comparison of chlonisol versus standard chemotherapy. *Notes:* Experimental –animals received chlonisol in MTD (20 mg/kg). Control – comparison group treated with lomustine or temazolomide in MTD.

In order to find a possible source of heterogeneity between studies, post-hoc analysis was attempted depending on the gender of the animals. In the subgroup analysis the Q-test showed statistically significant heterogeneity between studies in males and females (Chi^2^ = 2,81, df = 1; P = 0,09) and the I^2^ index was 64% (Figure 3).

**Figure 3.**
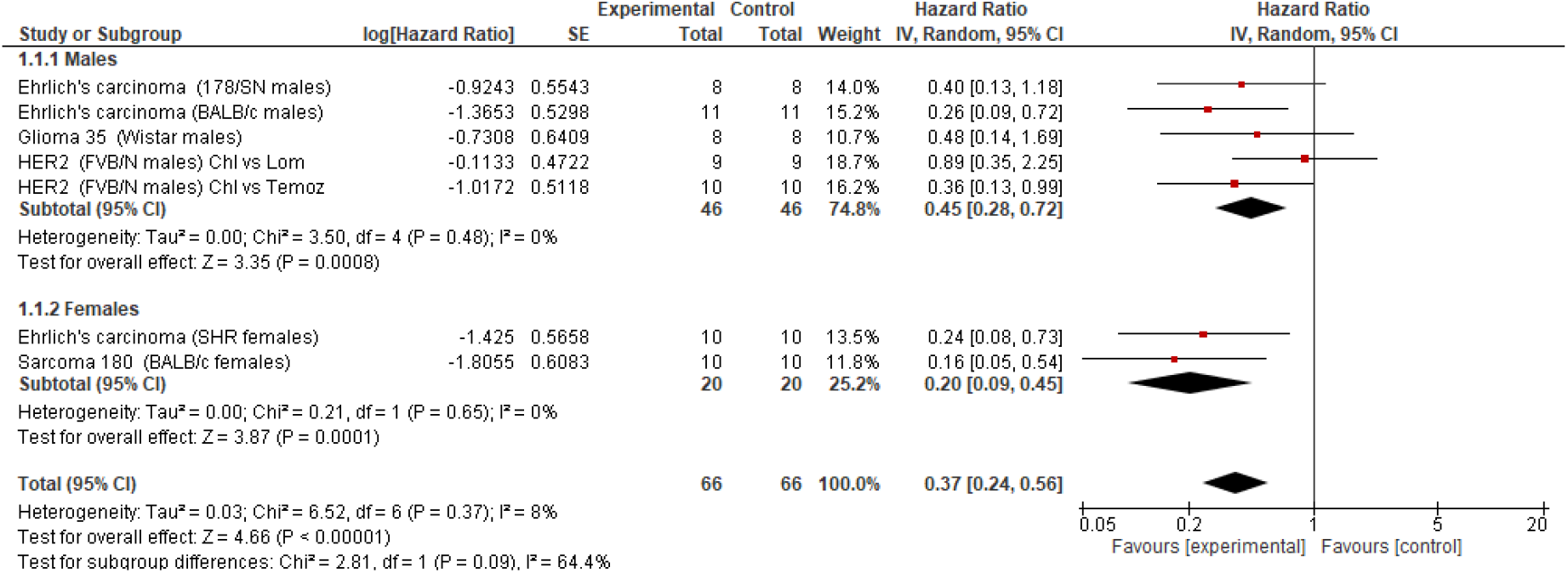
Random effects model for subgroup analysis of chlonisol versus standard chemotherapy in males and females. *Notes*: Experimental –animals received chlonisol in MTD (20 mg/kg), Control – comparison group treated with lomustine or temozolomide in MTD.

As an independent method of treatment, systemic chemotherapy for BT is rarely used because of its low ability to penetrate the blood-brain barrier (Jain 2018). The most commonly used chemotherapeutic drugs in the treatment of primary and metastatic BT are the imidazotetrazine derivative temozolomide and the NAU derivative lomustine. The monotherapy of these drugs has a low antitumor activity. Combining these chemotherapy drugs with surgical and radiation therapies has achieved some success (Zhu *et al*., 2014, Weller *et al*. 2020, Hotchkiss *et al*. 2021). Like lomustine, chlonisol belongs to the NAU derivatives. As specified by Ostrovskaya LA *et al*. (1998) for chlonisol, the ratio of areas under the concentration-time curves in brain and blood (AUCbrain/AUCblood) of animals with Lewis lung carcinoma was 10.5 when administered intravenously. This indicates that chlonisol is highly able to penetrate the blood brain barrier.

Meta-analysis offers a solution to the problem of synthesis of preclinical data by offering a robust statistical integration, powerful visualization and information collection tool, easy to interpret. Typically, meta-analysis procedure is preceded by publications searching with a predefined search strategy - systematic review. However, systematic reviews are not always necessary for preclinical studies. For example, if researchers have access to many quantitative research data. This situation is typical for pharmaceutical companies (Munoz-Muriedas 2021). Quantitative data for this analysis were obtained from our scientific laboratory’s internal preclinical report database. As a mathematical model, we selected a random effects model because it better reflects real situations, considering the nature and diversity of animal experiments (Hooijmans *et al*. 2014). Individual comparative preclinical studies of chlonisol in laboratory rodents, included in the analysis, demonstrated its effectiveness against a wide range of tumors transplanted intracranially. However, some studies have not achieved statistically significant effects of chlonisol compared with active control (see table 1). This may be due to the low number of rodents per group (8–9 animals). In this respect, the general statistical assessment of homogeneous preclinical studies appears to be well justified. A meta-analysis of preclinical studies showed that compared with control treatment, chlonisol administered in MTD of 20 mg/kg significantly reduced the risk of death of animals with intracranial tumors by 63% (HR=0.37; 95% CI: 0.24 -0.56; P <0.00001) (Fig. 2). Among the preclinical studies analyzed, a stable direction of the effect in favor of chlonisol was observed, only the size of its effect changed (HR and 95% CI). No significant between-study heterogeneity was observed (Tau^2^ = 0.03; Chi^2^ = 6.52; df = 6; P = 0.33; I^2^ = 8%).

The authors also sought to find sources of heterogeneity in the included studies. This attempt should not be viewed as data dredging, but rather as evidence of the possibility of subgroup analysis of preclinical studies using a specific example. In order to seek a possible source of heterogeneity between studies, *post-hoc* analysis was conducted depending on the gender of the animals. In the subgroup analysis the Q-test showed statistically significant heterogeneity between studies in males and females (Chi^2^ = 2,81, df = 1; P = 0,09) and the I^2^ index was 64% (Fig. 3). The low statistical power of the Q test should be taken into account due to the low number of studies in females. It should also be noted that comparison between males and females was not planned beforhead and the results should be interpreted with extreme caution. Nevertheless, this information can be used to create a hypothesis that chlonisol in women may have a more prominent antitumor activity.

## CONCLUSION

For the first time in the history of national experimental oncopharmacology, a meta-analysis of preclinical studies has been carried out. Using the example of domestic cytostatic, we presented a statistical synthesis of the results of individual studies to obtain a combined effect estimate of this compound in an animal model with a particular experimental oncological pathology. Analysis of the available preclinical studies showed that chlonisol administered in MTD of 20 mg/kg provides significant advantages in OS in laboratory rodents with intracranially transplanted tumors compared with lomustine and temozolomide. The results obtained can be used as a basis for further studies of chlonisol as a chemotherapeutic agent for primary and metastatic CNS tumors.

## Limitations of the study

1. Only the internal base of one laboratory was used.
2. Extremely low number of studies in females. The result of post-hoc subgroup analysis is hypothesis generating.
3. Lack of description of the method for calculating sample size in the studies.
4. Lack of detailed information on randomization technique, the presence of blinding groups of animals and information on the analysis of results obtained openly/blindly.

## REFERENCES

Bates A, Gonzalez-Viana E, Cruickshank G, Roques T & Guideline Committee (2018) Primary and metastatic brain tumours in adults: summary of NICE guidance. BMJ 362: k2924.

Jain KK (2018) A Critical Overview of Targeted Therapies for Glioblastoma. Front Oncol 8: 419.

Jones DTW, Banito A, Grünewald TGP, Haber M, Jäger N et al. (2019) Molecular characteristics and therapeutic vulnerabilities across paediatric solid tumours. Nat Rev Cancer 19(8): 420–438.

Hooijmans CR, IntHout J, Ritskes-Hoitinga M, Rovers MM (2014) Meta-Analyses of Animal Studies: An Introduction of a Valuable Instrument to Further Improve Healthcare. ILAR J 55(3): 418–426.

Hotchkiss KM, Sampson JH (2021) Temozolomide treatment outcomes and immunotherapy efficacy in brain tumor. J Neurooncol 151(1): 55–62.

Ioannidis JPA, Schmid CH, Lau J (2000) Meta-analysis in hematology and oncology. Hematol Oncol Clin North Am 14(4): 973–991.

Munoz-Muriedas J (2021). Large scale meta-analysis of preclinical toxicity data for target characterisation and hypotheses generation. PLOS ONE 16(6): e0252533.

Murazov IG, Stukov AN, Zmitrichenko IG, Niuganen AO, Tochilnikov GV et al. (2021) Evaluation of the antitumor activity of 2-[3-(2-chloroethyl)-3-nitrosoureido]-1,3-propanediol (chlonisol) in C57BL/6 mice with intracranially transplanted B16 melanoma. Pharmacokinetics and Pharmacodynamics 1: 23-29. (In Russ.).

Ostrom QT, Wright CH, Barnholtz-Sloan JS (2018) Brain metastases: epidemiology. Handb Clin Neurol 149: 27–42.

Ostrovskaya LA, Filov VA, Ivin BA, Stukov AN, Fomina MM et al. (1998) Chlonisol – the new alkylnitrosourea drug with antitumor activity. Russian biotherapeutic journal 3(1): 37–48. (In Russ.).

Stukov AN, Esikov KA, Usmanova LM, Kharitonova NN, Vershinina SF et al. (2020) Synthesis and Antitumor Activity of 2-[3-(2-Chloroethyl)-3-Nitrosoureido]-1,3-Propanediol (Chlonisol). Pharm Chem J 54: 579–581.

Vargo MM (2017) Brain Tumors and Metastases. Phys Med Rehabil Clin N Am 28(1): 115–141.

Vesterinen HM, Sena ES, Egan KJ, Hirst TC, Churolov L et al. (2014) Meta-analysis of data from animal studies: A practical guide. J Neurosci Methods 221: 92–102.

Weller M, Le Rhun E (2020) How did lomustine become standard of care in recurrent glioblastoma? Cancer Treat Rev 87: 102029.

Zhu W, Zhou L, Qian J-Q, Qiu T-Z, Shu Y-Q et al. (2014) Temozolomide for treatment of brain metastases: A review of 21 clinical trials. World J Clin Oncol 5(1): 19–27.

